# Unsuspected prevalent tree occurrences in high elevation sky-islands of the western Alps

**DOI:** 10.1101/2020.07.08.193300

**Authors:** Gilles André, Sébastien Lavergne, Christopher Carcaillet

## Abstract

The present note intends to challenge, based on field observations, the definition of treeline and argues for considering the upper occurrences of tree-species as an integrative result of historical and contemporary processes acting on high-altitude socio-ecosystems. A field survey of *Pinus cembra* growing above 2800 m asl was conducted, in an ecoregion of southwestern Alps, straddling France and Italy (Queyras-Mt Viso area). Pines were described (height, density) and their habitats were contextualized (altitude, slope degree and exposure, bedrock). Individuals of *Pinus cembra* are commonly growing between 2800 and 3200 m asl, generally on east-facing, steep and rocky slopes, with a pine density decreasing with altitude. Pines are generally dwarf-shaped, sometimes erected or prostrated. Their needles are half the size compared to those of pines growing in subalpine forests. Pine morphology indicates harsh growing conditions, notably strong wind, aridity and frost. These habitats constitute visually emerging objects attractive for the nutcracker caching behaviour. This setting acts as a positive passive driver for pine at high altitudes, which is further favoured by intense atmospheric moisture supplied by the east from the Po valley. Steep and rocky slopes exclude livestock, likely explaining why these pines are located in grass-less habitats. These tree clusters constitute remarkable “sky-island” populations. *Larix decidua* and *Pinus uncinata* were also recorded at exceptional altitudes but *P. cembra* is the most frequent species in high alpine conditions. These observations altogether highlight a complex pattern of tree-species line in the western Alps, and probably beyond.

## Introduction

Understanding the patterns and processes underlying upper location of trees, ie the so-called treeline altitude (TLA), has long been a topic of research and debates (eg, Tranquillini 1979; Dahl 1986; Holtmeier 1994). Unravelling the drivers of TLA has indeed far reaching implications to understanding vegetation dynamics under global changes, and TLA is notably used as a climate-proxy in paleoecology (Rochefort et al. 1994; Kullman 1995). However, a key question is whether TLA should be defined based on the physiognomy of tree species or on mere occurrences of trees. Indeed, many considered TLA when trees are erected and greater than few meters (eg, Payette 1983; Eronen and Huttunen 1993; Kaspar and Tremel 2018), growing in rather dense stands (cluster, tree-island; eg, Holtmeier and Broll 1992) or even visible by aerial or satellite imagery (Paulsen and Körner 2001; Gehrig-Fasel et al. 2007). We argue that it is important to distinguish the treeline and what we term the tree-species line, which line is the biological limit of a species whatever their physiognomy and the stand structure.

The physiognomy of trees directly depends on their very local environment (eg, micro-climate, topography, soil). During their individual life, dwarf trees can alternate between erected and prostrated stature, due to changing climate altering the winter survival of meristems and organs (Lavoie and Payette 1994), or possibly due to changes in herbivore pressure and in parasites outbreaks (Körner 1998). The tree-species line can thus constitute either the relict of TLA with ancient erected trees that have been stressed (eg, Laberge et al. 2001), or the forefront of a current tree expansion due to the release of aforementioned stressors (eg, Kullman 2001; Brodie et al. 2019). If the physiognomy of tree individuals is shaped by ecological processes, the mere presence of trees could be considered as a part of the treeline, notably in terms of potentialities of regeneration, biogeochemical assimilation for growth, and interactions with ecological partners. This is particularly true if high-elevation single tree occurrences have been largely overlooked and underestimated (eg, Oberg and Kullman 2011), which may be the case in the poorly accessible and harsh cliff environments often prevailing at higher altitudes (Dentant and Lavergne 2013).

TLA is certainly not a homogeneous ecological object. It varies tremendously according to species, and to local or regional contexts, thus resulting biogeographical peculiarities, which may be contingent to many historical processes, notably land use. For instance, high mountains in southern Europe have long been used for domestic livestock grazing, with great variability in space and time (Carrer 2015), often after forest clearing by means of fire (Leys and Carcaillet 2016). This suggests that the high-altitude occurrences of trees are the result of complex socio-ecosystemic processes. A given tree-species line is thus composed of isolated individuals that could be the relict of a denser treeline altered by centuries or millennia of livestock grazing and climatic pejoration, or the pioneers of the future TLA resulting from climate warming and land use abandonment.

In western Alps, the TLA is classically considered to occur near 2100-2200 m above sea level (hereafter “asl”) in northern massifs, and ca 2300-2400 m asl in the southern ones. TLA follows a clear gradient, being lower in colder and wetter massifs (i.e. northern and outer massifs, respectively) than in warmer and drier massifs (i.e. southern and internal ones). However, on the south-facing slopes of the Mount Viso, highest tree patches reach 2500 m asl (Motta and Nola 2001) suggesting a certain spatial variability. Trees at exceptional altitudes in different massifs of western Alps have been reported (André 2016; Carcaillet and Blarquez 2019). Here, we present a systematic survey of *Pinus cembra* L. occurrences located above 2800 m asl, from a French-Italian ecoregion, considering the frequency of individual number and height according to altitude, slope exposure and degree. This data contributes to change current viewpoint about the regional TLA of this pine species and others, as we report unexpectedly numerous isolated tree clusters growing at exceptional altitudes. This survey is supplemented by data collected more extensively in the western Alps. We then discuss these data in the light of known past and current environmental processes.

## Material and Methods

### Study species

*Pinus cembra* is a keystone tree species growing in subalpine forests of the Alps and the Carpathians. With *Larix decidua* Mill., *P. cembra* co-dominates forests of the Alps, where the continental-type climate is optimal for its growth. Populations in the peripheral mountain massifs are sporadic and sparse. *P. cembra* is a long-lived species (>500 years) with long generation intervals, which can maintain stable population size over long periods of time without inbreeding depression and preserve high amount of genetic variation (Gugerli et al. 2009; Toth et al. 2019). The species produces heavy seeds that are exclusively dispersed by a mutualistic bird (*Nucifraga caryocatactes* L., spotted nutcracker) specialized in extracting seeds of *P. cembra* from its non-opening cones and in caching seeds in soil far from the tree-mother (Crocq 1990). This ornithochory limits the dispersal distance of seeds from a few hundred meters from seed-source, and hinders the colonization potential of this pine, compared to the wind-dispersed *L. decidua* or *Pinus uncinata* Ram. ex. DC. *P. cembra* interacts with fungi, both symbiotic for nutrients and water acquisition and pathogens (Merges et al. 2018), These interactions would control its recruitment rate at high altitudes (Neuschulz et al. 2018).

### Data collection

Summer excursions in the western Alps have allowed to record occurrences of *Pinus cembra* from 2600 to 3200 m asl. The study area corresponds to inner valleys and massifs between 6.75°E-7.15°E and 44.60°N-44.78°N. This area covers, from west to east, the Queyras Massif notably the southern slopes and ridges of the Guil valley, and the Varaita valley with the Mount Viso Massif bordering the valley to the north (Fig. 1a).

**Figure 1.**
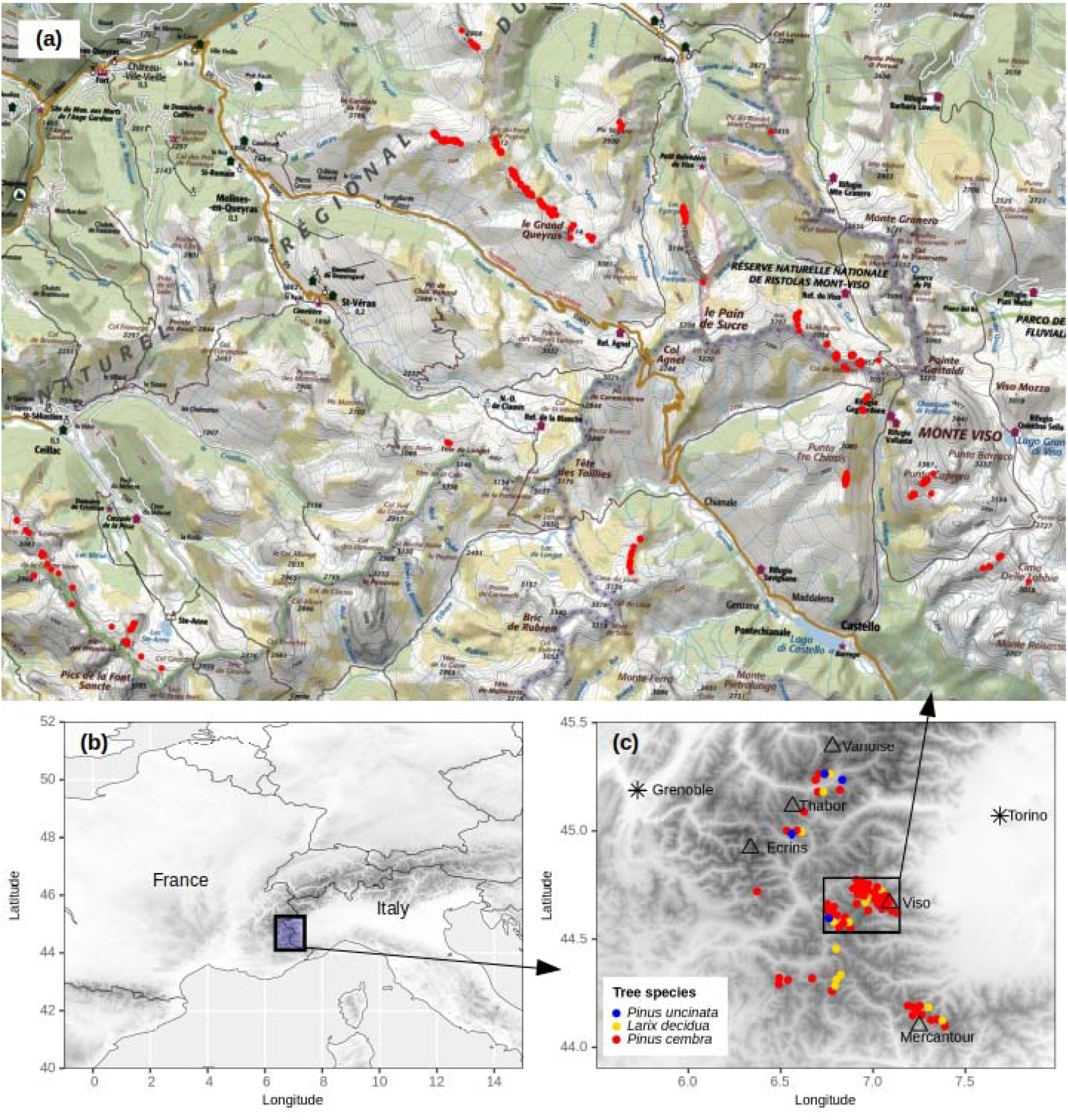
**(a)** Map of isolated *Pinus cembra* stands (red dots) in the area of the Queyras Massif and Mount Viso, southwestern French-Italian Alps, above 2800 m asl. On this map green areas correspond to forest whereas white and yellow areas correspond to grasslands and rocky outcrops (Map Data from Géoportail ©IGN-2020); **(b)** Location of the study area in the southwestern Alps straddling France and Italy; **(c)** Map of maximal altitudes of isolated trees (*Larix decidua, Pinus cembra, Pinus uncinata*) in the southwestern Alps; only individuals above the classic tree-line were recorded (these data are indicative); names of main massifs evoked in the text are indicated.

Hundreds of pine occurrences were mapped using GPS during excursions or positioned afterward using online maps and aerial photographs (geoportail.gouv.fr/). The heights of individuals were measured when possible or estimated (±10 cm) for trees inaccessible without alpinism equipment. Data were also collected to describe trees’ physiognomy, patch structure (isolated *versus clustered*), and ecological contexts (substratum, slope exposure and degree). Pictures were systematically taken to identify trees and contextualize their habitat and their growth forms.

We computed the percentiles of slope exposure of trees to explore the different patterns according to altitude, and latitude as a secondary factor. The slope exposures of high-altitude trees were compared to slope exposure of the 3100 field-data of *P. cembra* stands, extracted from the database of the National Alpine Botanical Conservatory (CBNA). Because the tree-density differed between the CBNA database and the our data, and also for the need of graphic representation of exposure of individual numbers, these densities were minimax transformed. Such transformation rescales the individual numbers from a given exposure class to range between 0 and 1 by subtracting the minimum numbers found in an altitudinal class, and dividing by the range of values of this altitudinal class.

### Additional data

Individual pines recorded in our study are small, maybe of high conservation interest, and therefore were not cored for tree-ring counting, although dendrochronology would certainly be of high scientific interest (eg, Mathaux et al. 2016; Büntgen et al. 2019). However, for testing purpose, one broken stem was collected on the ground at 2920 m asl on the Mt Viso, and one branch on an erected tree at ca 2500 m asl further north (45.181°N, 6.716°E) was cut at the branch origin at a height of 0.5 m. These both *P. cembra* samples provide indications on the potential growth and age of individuals. Whenever possible, the size of needles was noted, and the occurrences and physiognomy of cones were recorded.

## Results

### Altitude, topographic context, habitat, slope degree and exposure

A total of 544 *Pinus cembra* were recorded above 2800 m asl in the Queyras-Viso ecoregion. All tree-individuals reported here are situated above 2800 m asl, generally on ridges, often on steep and rocky habitats. Their main spatial distribution corresponds to a ridge oriented NW-SE including several main summits, namely the Grand Queyras, the Pain de Sucre and the Mount Viso (Fig. 1a). The surrounding valleys are the Aigue Agnel Valley (at northwest) and the Guil Valley (at north) both in France, and the Varaita Valley (at southeast) in Italy. Secondary stands are situated in the northwestern part of the Varaita Valley near the Longet Pass that connects with the Ubaye valley in the west, and the Crête des Veyres, which is a ridge oriented NW-SE between the Ceillac Valley and the Escreins Valley both in France (Fig. 1a).

The frequency of individuals decreases with site elevation, reaching the exceptional altitude of 3200 m (Fig. 2a). The median altitude of observations is 2835 m (mean ±sd: 2853 ±58 m asl). A proportion of 75% of inventoried pines occurred below 2870 m asl, suggesting that pines observed above 2870 m have a very exceptional altitudinal location. The frequency distribution of pines along altitude classes reveals a negative exponential (r^2^=0.99; Fig 2a) corresponding to a halving of individuals at every ~40 m of elevation difference.

**Figure 2.**
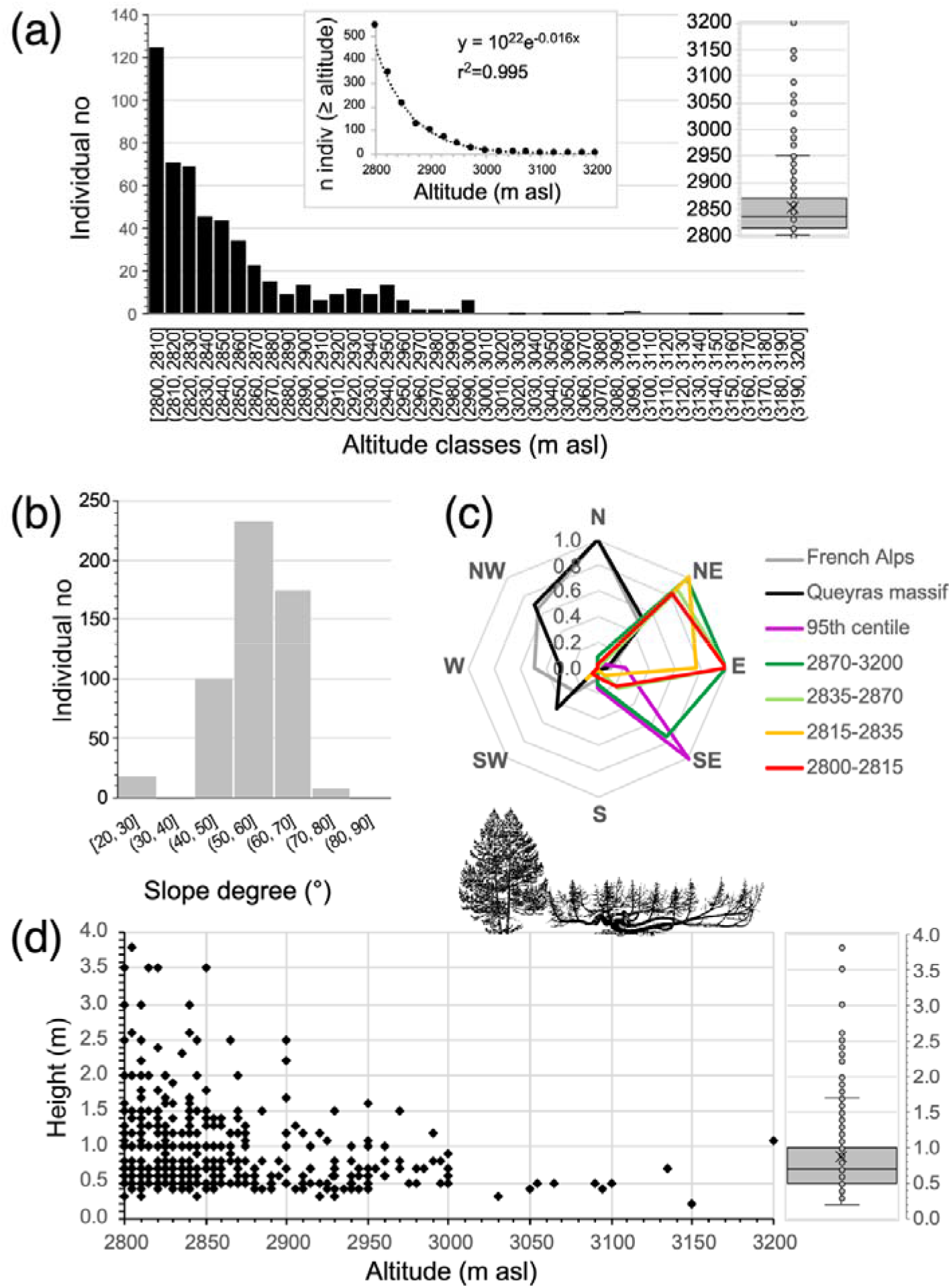
Distribution of *Pinus cembra* according to **(a)** altitude from 2800 to 3200 m asl (in insert: box-plot distribution of altitude of trees), **(b)** the slope degree, and **(c)** the slope exposure in the Queyras-Viso ecoregion. The distribution of exposure **(c)** of isolated *P. cembra* (n = 544) are sorted per altitudinal classes, and compared to the values for *P. cembra* stands mostly in forests of the Queyras massif and from the French Alps (n = 1747); numbers were minimax-rescaled to compensate for differences in observation numbers; **(d)** tree-height of *Pinus cembra* plotted against altitude; the median height is 0.7 m, and the 75^th^ percentile is 1.0 m (in insert: box-plot distribution of heights).

These pines are located on cliffs and rocky outcrops, generally with steep slopes. The modal distribution of slopes is 50-60°. About 94% of all individuals were found on slopes between 40° and 70° (n = 511; Fig. 2b). Some pines, although quite rarely, grew with grasses but always embedded between rocks or within rocky outcrops. On cliff and ridges, individuals are rooted in cliff cracks or between bedrock layers offering a softer and weathered material (Fig. 3a-c).

**Figure 3.**
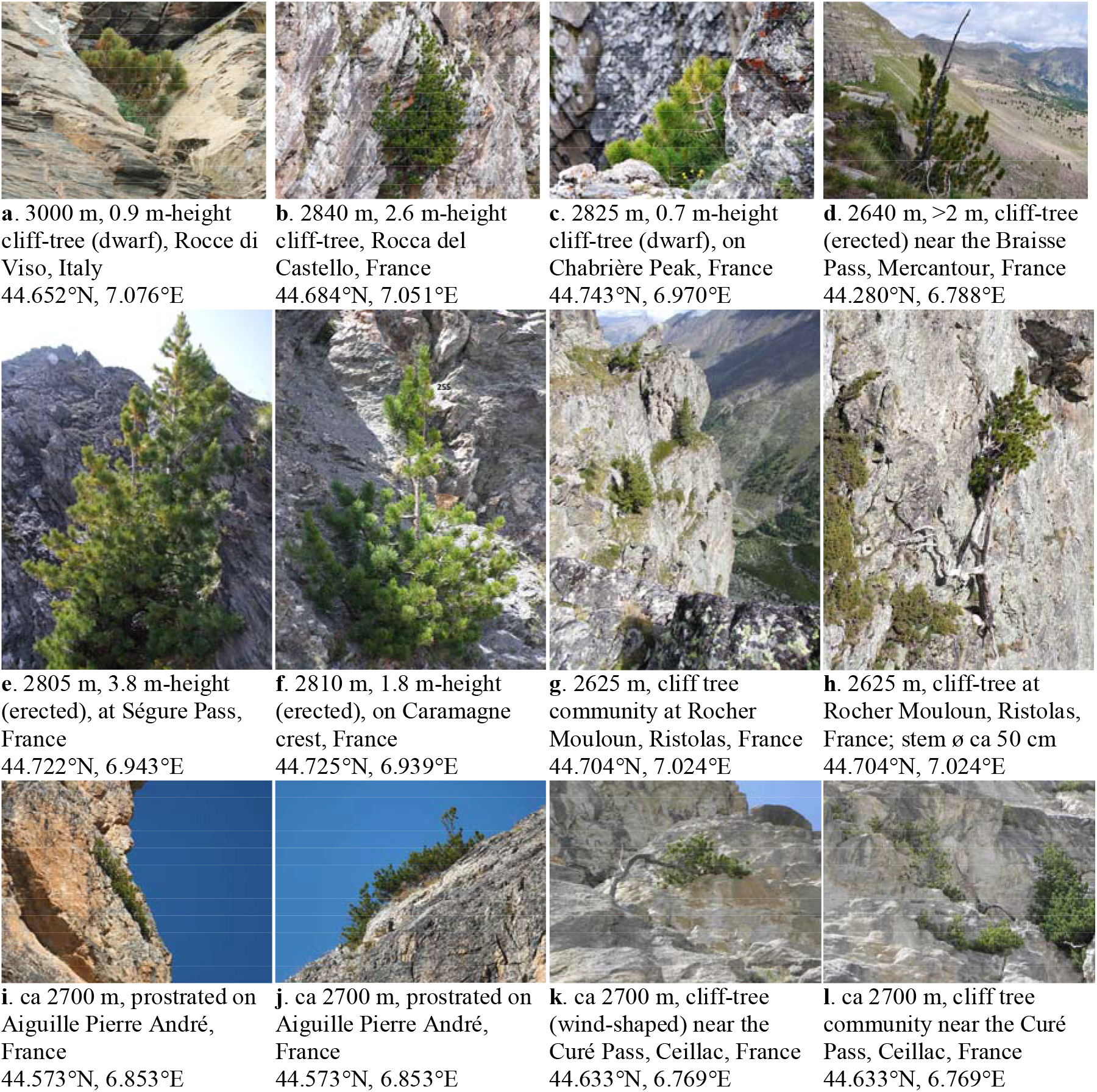
Examples of *Pinus cembra* growth form between 2600 and 3200 m asl in the western Alps, in France and Italy. These pictures display different contexts of growth (cliff, outcrop) on different chemical types of rocks (eg, ophiolite [g,h], quartzite [i,j], limestone [k,l], calcareous sandstone [d]), and of tree physiognomy (prostrated, dwarf, erected, flag-shaped). All pictures are from GA, except the [d] from CC.

The exposure of individuals shows that, on average, pines located above 2800 m generally occur on east-exposed slopes (main mode), ranging from NE to SE aspects (Fig. 2c). Interestingly, when the individuals are sorted by altitude and latitude, and clustered per percentiles, the pines located below 2870 m (75^th^ percentile) all occur on slopes facing east to northeast. The upper quartile of individuals situated between 2870 to 3200 m asl shows a similar pattern but with slopes more often facing southeast. When considering only 95^th^ percentile (ie, n = 27, alt. ≥ 2960 m asl), they show a general exposure to the southeast. These patterns of slope exposure of highest records of *P. cembra* strikingly contrast with the common exposure of subalpine forest stands in Queyras and French Alps, which mostly occur on north-facing slopes (Fig. 2c).

### Individual physiognomy, density and spatial pattern, biological singularities

Censused pines are generally growing in clusters consisting a few up to tens of individuals. Their density can attain 20 individuals per hectare. These pines are generally growing in disconnected stands from subalpine forests situated in the valley below 2200-2400 m asl, sometimes at hundreds or thousands of meters from nearest woodlands (Fig. 1a). However, some stands are connected to forest by the way of a narrow corridor of trees growing among rocks or on rocky ridges from the forest to these isolated stands. Spatial pattern of patch isolation and spatial structure was thus not homogeneous across censused trees and patches.

Most trees are dwarf-shaped (Fig. 3a-c), while very few are prostrated (Fig. 3ij), or erected (Fig. 3d-g). Flag-shaped trees were observed below 2800 m (Fig. 3hk). The median height is 0.7 m ranging from 0.2 to 3.8 m (Fig. 1d). Stem diameters are generally proportional to their height, but their growth rate is very low according to the two test samples realized. The broken stem collected at 2920 m presents about 45 tree-rings of about 250 μm-thick in average while the cut branch shows about 70 tree-rings of ca. 120 μm-thick in average. Female cones of *P. cembra* have been observed on few individuals up to 2855 m asl, but these cones were small and malformed. Needles of these pines were smaller (down to 4 cm) than needles of pine trees growing in larger forest stands at subalpine elevations (10-12 cm).

## Discussion

These observations demonstrate that *Pinus cembra* can often grow at exceptional altitudes as high as 3200 m asl in the western Alps. Although studies previously reported pines growing way in high altitude (Bono and Barbero 1971; Paulsen et al. 2000; Motta and Nola 2001; Carcaillet and Blarquez 2019), they never depicted such exceptional figure of altitudes for the Alps, nor such local abundance. These scattered populations of isolated *P. cembra* are often geographically disconnected from the main altitudinal range of trees situated sometimes 400 or 600 m lower in elevation. The very peculiar topographic situation of these tree occurrences recalls us the one of trees growing on isolated Madrean mountain ranges (Arizona), which were popularized under the metaphor of “sky-island” (Heald 1967). Below we discuss the ecological context of these sky-island trees, and put their particular habitats into perspective with history of land use practices for animal alpine husbandry. Last, these observations are discussed in terms of species-line *versus* classic treeline notably making analogies with the limits of tree populations during glacial periods.

### “Premier de cordée”: frugal and dwarf alpinist trees

These isolated and elevated trees exclusively occur on rocky outcrops, steep slopes and ridges. Such inhospitable habitat was unexpected at a first sight, and raises questions about plant nutrition. Indeed, in rupicolous conditions, there are no or little water storage capacity. The humidity depends on the precipitation which supplies water to roots, but water are rapidly drained due to gravity. The important cloudiness on summits also supplies moisture in the form of vapor, because the area receives humid warm airmasses coming westward from the Po River plain and the Adriatic Sea, notably during growing season. Further, at high altitudes, the chemical weathering processes are more limited than at lower elevations due to low temperatures (Riebe et al. 2004), and the steep slopes favour the physical erosion of elements and particles (Larsen et al. 2014). These conditions thus result in a low soil rate formation and limited nutritional capacity. The high-altitude trees certainly have a particular nutritional physiology implying rates and assimilation closely linked to the ectomycorrhizal capacity of seedlings that seems not limited by the altitude (Merges et al. 2018). Generally, ectomycorrhiza are crucial for trees to capture soil humidity and to recycle nitrogen from organic matter (Smith and Read 2008). The nutritive capacity of high-altitudes trees is probably rather different than that of trees growing in the subalpine forest or even at the classic TLA, thus impeding any comparison with the biology of the low-altitude trees.

Site elevation also appears to be a strong driver of these pine occurrences. The negative exponential modelling of pine density according to altitude (Fig. 2a) indicates that the pine survival is a function of altitude, probably regulated by a mortality or recruitment constant (Hett and Loucks 1976). The dispersal function of *P. cembra* seeds depends on nutcrackers, which preferentially caches seeds in soil of closed dry forests, but the high probability of seed caching is negatively correlated to seedling establishment in habitat of quality (tree-less habitat, wet soil; Neuschulz et al. 2015). The very rare stored seeds in soil caches of wet and open habitat have thus a maximum probability of efficient recruitment. Above 2800 m, the habitat is dry and far from seed-source, and thus *a priori* unfavourable for seedlings especially on south-facing slopes, which are submitted to severe evapotranspiration due too strong solar irradiance. However, east-facing slopes where 95% of pines are located (Fig. 2c) are more humid thanks to lower irradiance, all other factors being equal. East-facing slopes with low solar irradiance would thus compensate for the aridity of south-facing slopes. Above 2970 m, most pines grow in south-facing slopes. This particular exposure might compensate for shorter growing season with increasing elevation by higher temperatures (Fig. 2c).

The growth forms and physiognomies of these high-altitude pines mostly correspond to prostrated and dwarf statures, but also erected trees, sometime damaged by wind, snow or lightning (Fig. 3). These local peculiarities of high elevation tree occurrences may challenge the conventional vision of the upper treeline, and question the reason to consider only erected trees to determine TLA. The observed decrease of tree height with elevation (Fig. 2d) might not necessarily be due to a younger age of highest individuals resulting from recent recruitment. Similar observations of decreasing pine height with elevation showed a reduced length of annual shoots in high altitude environments prone to strong wind and low temperatures (Takahashi and Yoshida 2009). We also reported that needles of *P. cembra* above 2800 m are smaller than those from subalpine forest trees (ca 4 cm vs 10-12 cm). Altogether, the observed reduced leaf length, shoot length and individual height could as well be morphological plasticity or even adaptation to the harshness of high altitude (eg, Schoettle 1990; Kajimoto 1993). The dwarf and prostrated physiognomy of high altitudes trees should be considered as the result of a harsh environment rather than function of time since regeneration. Such isolated high-altitude trees may thus be considered as the real fundamental niche limits of *P. cembra*. More investigations into these sites’ microclimate and relation to the known fundamental niche of lower elevation forest stands would be very interesting for better understanding niche and range dynamics of these cold-climate cembra pines.

### The unbearable doubtness of land use

The localisation of these elevated rocky habitats of isolated trees where grasslands are absent or very scarce strongly argues for involving livestock grazing as the chief mechanism excluding tree-species from alpine grasslands. Livestock acts through grazing and trampling of seedlings and browsing of saplings and dwarf trees especially *Larix*. Wild herbivores would be not involved, or only marginally, because they have naturally low-density populations compared to domestic livestock.

The deforestation time above 2600 m for the needs of husbandry is difficult to accurately assess. Fire history can be used as a proxy of land use expansion based on the fact that people used fire as an easy process to reduce the woody cover in favour of grasslands for livestock dedicated for dairy production based on cattle (Carrer 2015). Interestingly, a sedimentary charcoal study at 2214 m asl in the Queyras Massif has revealed a maximum of fire frequency between 4000-3500 cal yr BP, corresponding to a progressive increase from 2 to 4 fires per 1000 years (Carcaillet and Blarquez 2017). This trend matches soil charcoal-based studies in the same massif reporting fires with an increase of their occurrences after 4000 cal yr BP (Saulnier et al 2015). However, all these fire proofs are situated below 2800 m asl. Above 2400 m, soil sequestrated charcoals are so rare and small that they limit the implementation of paleo-studies (Carcaillet and Brun 2000; Talon 2010). The rise in prehistorical social activities in high altitude is dated of 4500 to 4000 cal yr BP in the western Alps according to the changes and the intersites heterogeneity of fire frequencies (Leys and Carcaillet 2016).

### “Sky-island” trees: from treeline to tree-species line

Some pines are growing up to 800 m above the classic TLA. These altitudes are definitely altitudinal records of tree occurrences for the Alps, maybe facilitated by the rather southern location of the Queyras and Viso massifs compared to northern (eg, Vanoise) or western massif (eg, Écrins), which are wetter and colder. However, few other sites in the north and the south of western Alps also present high-altitude trees attaining 2700 (*Pinus uncinata*) to 2800 m (*Larix decidua*, *P. cembra*; Fig. 1c), but none presents such records with trees 800 m above the local TLA. Interestingly, these regional peculiarities are situated on extremely heterogeneous bedrocks, with individuals on dolomitic limestone (Fig. 3kl), calcareous sandstone (Fig. 3d), calcschist, flysch, ophiolites (Fig. 3gh), basalt, quartzite (Fig. 3ij), micaschist, or acidic schist and sandstone, which indicates that the chemical quality of bedrock is likely not a significant driver or limitation for the sky-island trees.

A magnitude of 600-800 m between the species-line altitude and the classic TLA echoes to the finding of tree remains, mostly needles of *P. cembra* and few of *L. decidua*, in lacustrine sediments dating from the end of the last glaciation assuming the occurrence of glacial refugia on a *nunatak* in the Queyras massif, ie an area uncover by glaciers (Carcaillet and Blarquez 2017). The lake is currently situated at 2214 m asl, ie about 50-100 m above the trimline of the last glaciation (Cossart et al. 2012). If this finding corresponds to the species-line, the TLA would have been situated 600-800 m lower, ie at 1400-1600 m. Because the valleys were filled-in by glaciers with emerging *nunataks* (Buoncristiani and Campy 2011; Cossart et al. 2012; Seguinot et al. 2018), the nearest available areas around 1400-1600 m asl in front of the glaciers were in the east in the Italian foot-hills, or south near the current French city of Sisteron where fossil trunks of *Pinus* cf. *sylvestris* have been found (Miramont et al. 2000) making this area a peripheral glacial refugium (*sensu* Holderegger and Thiel-Egenter 2009). If Carcaillet and Blarquez (2017) evokes once the treeline, their finding mostly refers to the tree-species line, which does not involve erected trees. Based on the present study, we can figure dwarf or prostrated pines or larches, similar to individuals reported here, growing on *nunataks* in the glacial refugia of the Queyras massif.

It is possible that inventories in other sites, notably in the Alpi Marittime European Park straddling France and Italy in the south (cf. Mercantour; Fig. 1c), and further north around the Thabor Massif (Fig. 1c), could provide similar patterns of “sky-island” trees as in the Queyras-Viso ecoregion. In addition, it has to be considered that such as higher occurrences of isolated trees have largely been overlooked, due to a lack of botanical prospections in high alpine and nival belts (eg, Körner et al. 2011; Dentant and Lavergne 2013).

## Acknowledgements

Authors are indebted to the IGN for the authorization to reproduce the map (Fig. 1a), and M. Luc Garraud of the CBNA for his help with the CBNA plant database.

## Compliance with ethical standards

### Authors contribution

AG carried out the fieldwork and collected most of the data. CC produced a draught of manuscript, and all co-authors contributed to further versions. All authors approved the final version.

